# Canaries differentially modulate solo and overlapping singing during the transition to the breeding season

**DOI:** 10.1101/2025.01.09.632199

**Authors:** Santhosh Totiger, Pepe Alcami

## Abstract

Songbirds sing different song types depending on the social context. Songs can be categorized into two types based on their timing relative to other birds’ songs: solo and temporally-overlapping songs. Overlapping songs have been typically characterized in the breeding season, in which they are associated with an aggressive social context. However, whether both song types occur year long, and whether they show differential modulation of their frequency and properties during the transition between the non-breeding and the breeding season has been rarely studied. Here we investigate, in a group of domesticated canaries *(Serinus canaria)*, the variation of singing in both song types at the transition between the nonbreeding and the breeding season. We found that both song types were present outside the breeding season. Whereas for solo songs, duration and its variability, fraction of time singing and number of songs increased as the breeding season approached, overlapping songs showed opposite trends. Furthermore, both song types were distributed more homogeneously as the daylength increased. Overall, the differential changes of both song types at the seasonal time scale suggest differential underlying mechanisms and functions of these two song types found in a social setting.

## 1. Introduction

The evolution of animal behaviour, including acoustic communication, has been shaped by seasonal changes (1). Seasonality of animal communication has been observed in a number of taxa, some animals producing more acoustic signals at a particular time of the year, for example frogs (2, 3), birds (4), mammals (5) and primates (6). Among tetrapods, songbirds produce elaborate vocalisations called songs in the context of territorial defence, mate guarding, reproduction, and social communication (7). Birdsong can be categorised into different types depending on the social context. One such categorisation is based on the temporal coincidence of two or more birds singing during a fraction of their song. These songs are referred to here as ‘overlapping songs’, and individual bird singing is referred to as solo songs. These song types are acoustically and functionally different, overlapping singing having been reported to carry aggressive intent in the breeding season (8, 9). Although a number of studies describe seasonal variation in singing behaviour (10-14), how these song types vary at the transition between the non-breeding and the breeding seasons in a social setting remains elusive.

Canaries (*Serinus canaria*) are a seasonally-breeding songbird species. Their songs undergo seasonal plasticity in duration, acoustic and syntactic properties (15-18). As a result, canaries have emerged as a model species to study adult behavioural plasticity and its underlying neuronal plasticity (19). In this context, previous studies have focused on solo songs. However, canaries also sing temporally overlapping songs in the breeding season, which are accompanied by aggression (8). It is not known whether overlapping songs are present outside the breeding season. Moreover, whether solo and overlapping songs change during the transition between the nonbreeding and the breeding season is not known. In this study, in a group of domestic canaries in an aviary, we characterise (a) solo and overlapping songs outside the breeding season, (b) to what extent both song types vary at the transition between the nonbreeding and the breeding seasons.

## 2. Methods

### (a) Birds and song recording

The study was conducted in an outside aviary located at the Biocenter, Ludwig-Maximilians-University, Munich, Germany (48°06’32.1”N 11°27’25.6”E). The breeding and keeping of the birds was approved by the veterinary authority of the Munich District Office (file number: 4.3.2-5682/LMU/Department Biology II; dated 16/09/2019) in accordance with the German Animal Welfare Act and the Directive 2010/63/EU of the European Parliament and of the Council of September 22, 2010 on the protection of animals used for scientific purpose.

The aviary consisted of 22 canaries (15 males and 7 females). This is consistent with a larger proportion of males compared to females observed in the wild (16). During the recording period (21 December 2019 to 05 March 2020) one male died on 14 January 2020, that is, after 4 days since the first day of recording. From then on, and until the end of the study, the aviary consisted of 14 males and 7 females. Breeding starts in April in the aviary. The 11.4 m^2^ aviary included a wooden chamber where birds could shelter. The lighting condition was natural, and the room temperature was controlled with an automatic heating system ensuring that it did not fall below 15° C. Fresh food and water were provided *ad libitum*.

Audio recordings were performed for 18 days between the above-mentioned dates, from sunrise to sunset, using a Marantz Solid State Recorder PMD661 MKIII with built-in mono microphone at a 48 kHz sampling rate. The recordings were conducted from outside the aviary to minimize the impact of human presence on singing activity.

### (b) Song annotation and analysis

All songs were manually annotated by careful visual inspection of the spectrogram using Audacity software version 3.4.0 with a frame rate of 22050 Hz. We noted down date and time of singing, song duration, and type of song - either overlapping song or solo song. The entire data was annotated two times to ensure the annotation errors are minimized. We used a minimum song duration criterium of 1s and a minimum time gap of 0.5s to separate individual solo songs. If individual solo songs could not be differentiated due to the temporal coincidence of two or more birds singing, we considered the entire duration of the singing interaction as duration of ‘overlapping song’. In total, we annotated 9475 solo songs and 4661 overlapping songs.

We then calculated for solo and overlapping songs (i) the fraction of time singing, defined as the total singing duration divided by the total recorded time; (ii) the mean song duration; (iii) the total number of songs divided by the total recording time; the variation of song duration by calculating the (iv) the standard deviation (SD) of song duration; and (v) the coefficient of variation (CV) of song duration; (vi) the CV of the duration of silent intervals between songs to characterise the distribution of songs throughout the day. We analysed how these parameters vary with increasing daylength for both song types.

### (c) Statistical analysis

The statistical analysis was performed using R software version 4.4.1. (20). The seasonal data was fitted with a linear model using the base command “lm” and the “ggplot2” package for creating plots.

## 3. Results

### Solo songs seasonal variation

We observed an increase in mean song duration as the breeding season approached (Pearson correlation, R = 0.61, p = 0.007, figure 1 B, C1). Song duration became more variable as the daylength increased (SD, R = 0.73, p < 0.001, figure 1 B, C2), and this increasing variability in song duration remained when computing the coefficient of variation (CV) (CV, R = 0.68, p = 0.002, figure 1, C3). In addition, we observed an increase in the fraction of time singing (R = 0.82, p < 0.001, figure 1 D), and the number of songs per hour (R = 0.80, p < 0.001, figure 1 E) as daylength increased. Furthermore, we analyzed how solo songs are distributed in time computing the CV of time intervals between solo songs. We found that songs were more homogeneously distributed as the breeding season approached (inter-solo interval CV: R = -0.79, p < 0.001, figure 1 A2, A3 and F).

**Figure 1:**
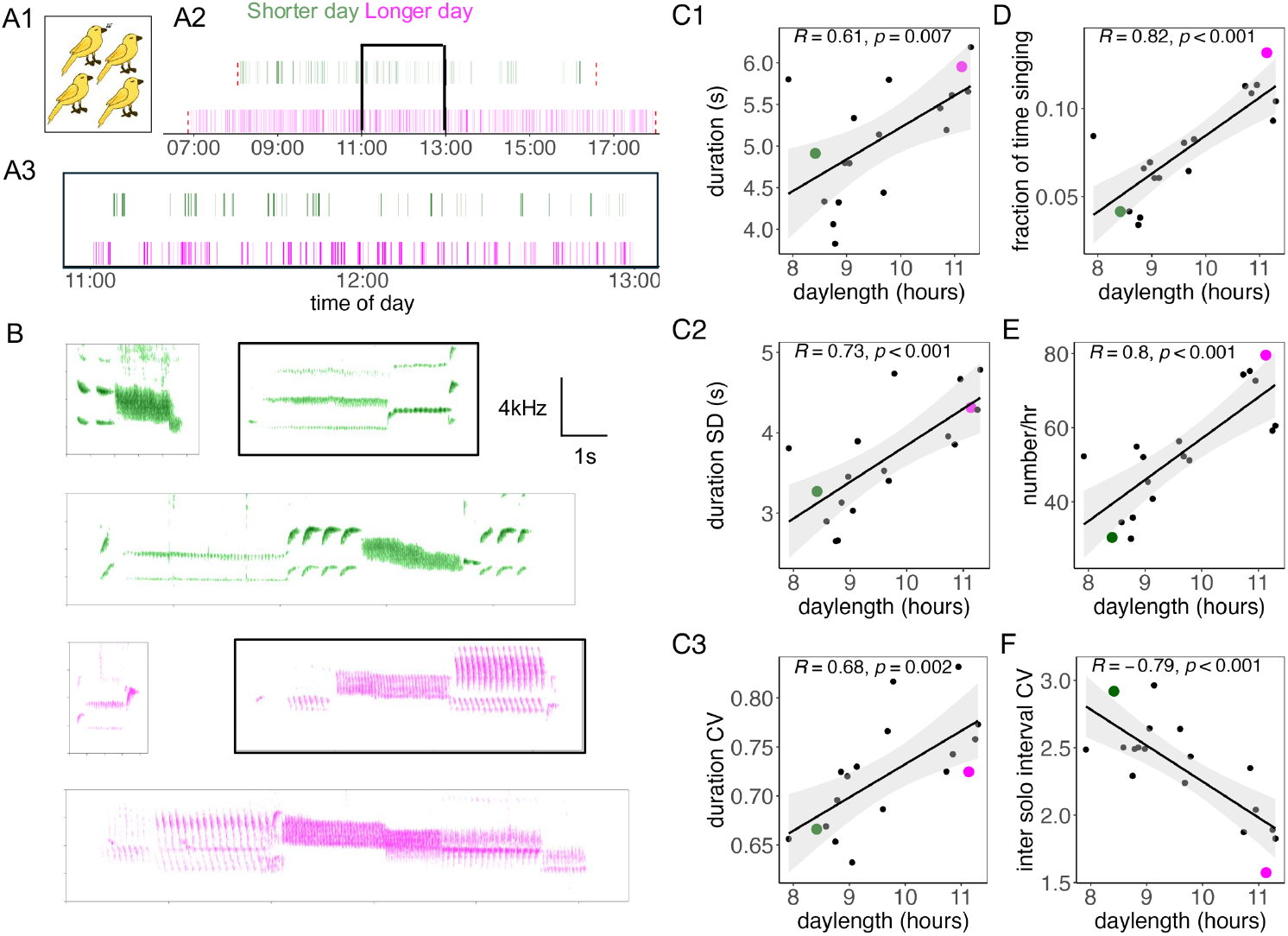
Seasonal variation in solo songs in a group of canaries. **A1**. Representation of solo songs, where one bird is singing, and the rest is silent. **A2**. Example of songs in representative days for solo song events from sunrise to sunset for a shorter day on 06 January (green) and a longer day on 02 March (magenta). Each vertical bar represents a song event. Red dashed lines indicate sunrise and sunset time. **A3**. Enlarged inset from the box in A2 shows the singing events in a two-hour time window revealing that songs are more numerous in the longer day. **B**. Example songs for a shorter (green) and a longer day (magenta); songs representative of mean-SD, mean (in black box), and mean+SD. **(C-F)** Measured parameters as a function of daylength: **C1**. Mean song duration. **C2**. Standard deviation (SD) of song duration. **C3**. Coefficient of variation (CV) of song duration. **D**. Fraction of time singing. **E**. Number of solo songs per hour. **F**. Inter-solo song interval CV. Each dot represents a day, examples from panel A1, A2 and B are taken from the days represented by enlarged green and magenta dots in C1-F. In C1-F the black line represents the linear regression fit and the grey area the 95% confidence interval. R; Pearson correlation. p; linear regression model p-value.

### Overlapping songs seasonal variation

The duration of overlapping interactions decreased as the breeding season approached (R = -0.75, p < 0.001, figure 2 B, C1). The variability of overlapping interaction duration decreased with increasing daylength (SD, R = -0.77, p < 0.001, figure 2 B, C2), and this decreasing trend remained after we computed the CV of song duration (CV, R = -0.76, p < 0.001, figure 2 C3). In contrast to solo songs, overlapping singing was more prevalent during shorter days and fraction of time singing decreased as the daylength became longer (R = -0.49, p = 0.038, figure 2 D). However, we observed no change as a function of daylength in their number per hour (R = -0.25, p = 0.312, figure 2 E). In addition, similar to solo songs, overlapping interactions were more homogenously distributed as the breeding season approached (inter-overlap interval CV: R = -0.79, p < 0.001, figure 2 A2, A3 and F).

**Figure 2:**
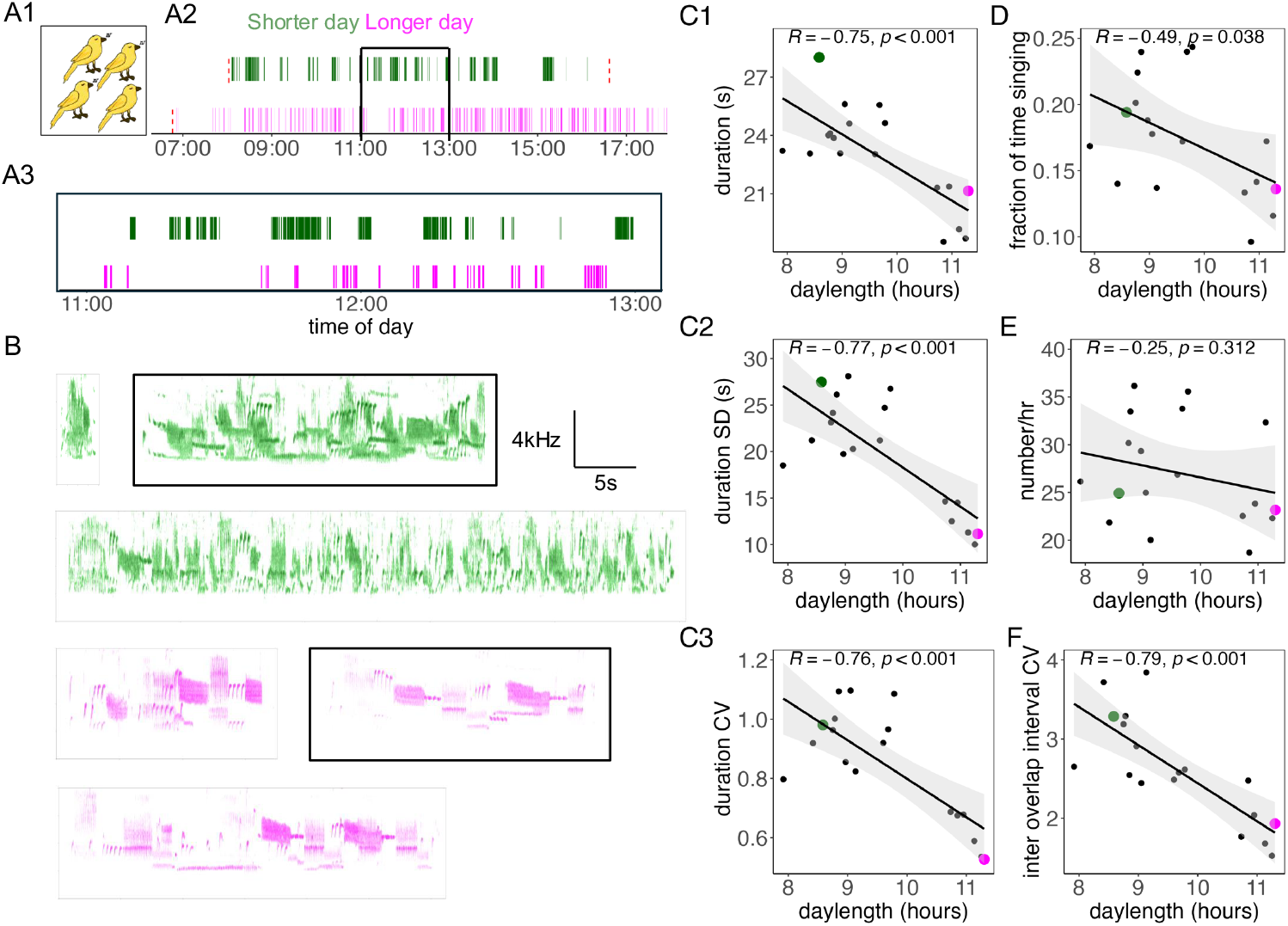
Seasonal variation in overlapping songs in a group of canaries. **A1**. Representative bird songs overlapping in time. **A2**. Example of songs in representative days for the overlapping interaction events from sunrise to sunset for a shorter day on 08 January (green) and a longer day on 05 March (magenta). Each vertical bar represents overlapping song events. Red dashed lines indicate sunrise and sunset time. **A3**. Enlarged inset from box in A2 shows the overlapping singing events in a two-hour time window. **B**. Example songs for a shorter (green) and a longer day (magenta); songs representative of mean-SD, mean (in black box), and mean+SD. **(C-F)** Following parameters as a function of daylength: **C1**. Mean song duration. **C2**. Standard deviation (SD) of song duration. **C3**. Coefficient of variation (CV) of song duration. **D**. Fraction of time singing. **E**. Number of solo songs per hour. **F**. Inter-overlapping song interval CV. Each dot represents a day. Examples from panels A1, A2 and B are taken from the days represented by enlarged green and magenta dots in C1-F. In C1-F, the black line represents the linear regression fit and the grey area, the 95% confidence interval. R; Pearson correlation. p; linear regression model p-value.

### Daily variation

Along with seasonal variation, we also analyzed daily variation in singing for both song types. We divided each day into three equal parts and calculated how the singing parameters change within a day. For solo songs, we did not observe any significant daily variation in fraction of time singing, duration, SD of duration, CV of duration, and number per hour (Wilcoxon rank-sum test, with Bonferroni-Holm correction) (Figure S1). For overlapping songs, we only observed daily variation in the fraction of time singing, lowest in the late period compared to the earlier period, and in duration which was shorter in the early period compared to the mid period (Wilcoxon rank-sum test, with Bonferroni-Holm correction) (Figure S2).

## 4. Discussion

It is well established that songbird songs, including canary songs, are seasonally plastic. Here we further characterise seasonal variation of two song types: solo and overlapping songs. Canaries solo song duration increased gradually and linearly as the breeding season approached. The magnitude of this increase is comparable to differences between the breeding and the non-breeding seasons reported in previous studies (15, 18). One key finding is that canaries spend more time singing overlapping songs, which were longer and more variable in duration, in the shorter days. In contrast, solo songs were sung more frequently and were longer and more variable in duration as the days became longer. In line with these results, white browed weavers and orioles sing duets more frequently relative to solo songs in the nonbreeding season, suggesting that differential seasonal modulation of solo and non-solo song types may be a general feature (21, 22). This opens the question of what the motivation and function of overlapping songs in the nonbreeding season is. We postulate that birds may practice and adjust while singing characteristic variable nonbreeding season songs in flocks before songs crystallise into the breeding season songs (15). Alternatively, an explanation for less overlapping interactions as the breeding season approached could be that males engage in longer overlapping interactions in the context of mate attraction in the non-breeding season and form pair bond. By contrast, overlapping singing may be avoided in a nonengagement manner in the breeding season (23). In the future, these hypotheses should be further investigated to unveil the function of overlapping songs in the nonbreeding season.

Our classification of songs into solo and overlapping here is instrumental and makes no assumptions about the intention of overlapping songs (9, 24, 25). However, if overlapping songs were the mere result of random overlaps of solo songs, we would expect similar variations as those shown by solo songs across seasons. Our data instead indicates that from the non-breeding to the breeding season, singing duration and fraction of time singing decrease for overlapping songs, whereas they increase for solo songs. This suggests that they constitute differentially-modulated song types. Moreover, it has been shown in canaries that the distribution of overlapping songs is not random and is accompanied by aggression in the breeding season (8). Although the aviary setting in the current study allows us to track the same population over time in a constrained space, the study has limitations. We took the approach to measure singing activity in a group of birds. However, individual bird contributions to driving singing dynamics across seasons remains unstudied. This should be addressed in the future using telemetry techniques such as backpack recordings in a wild population.

We show that not only solo songs undergo seasonal plasticity but also overlapping songs. Moreover, these song types change with contrasting trends in their plasticity. Underlying mechanisms could be seasonal changes in hormones and in brain circuits involved in singing. One candidate hormone is testosterone, whose levels rise as the breeding season approaches and is related to longer and more stable songs (26). However, whether and how these hormones are modulated during social interactions in relation to song type remains to be explored. At the neuronal level, circuits involved in the differential plasticity of solo vs overlapping songs across seasons require further investigation (15). In summary, the results in this study bring new questions on the mechanisms underlying song plasticity and adult brain plasticity in a social context.

## Acknowledgements

We are grateful to Dieter Leippert, and the animal caretakers for the excellent care and monitoring of the birds, especially to Sabrina Schenk. We are thankful to Prof. Benedikt Grothe for support, to Prof. Manfred Gahr for feedback on the project, and to Stefan Leitner for the comments on a previous version of the manuscript. The work was supported by LMU Biomentoring program of the faculty of Biology at the LMU (PA), Lehre@LMU program (ST) and a scholarship from the Studienstiftung des deutschen Volkes (ST).

---

**Supplementary figure S1.**
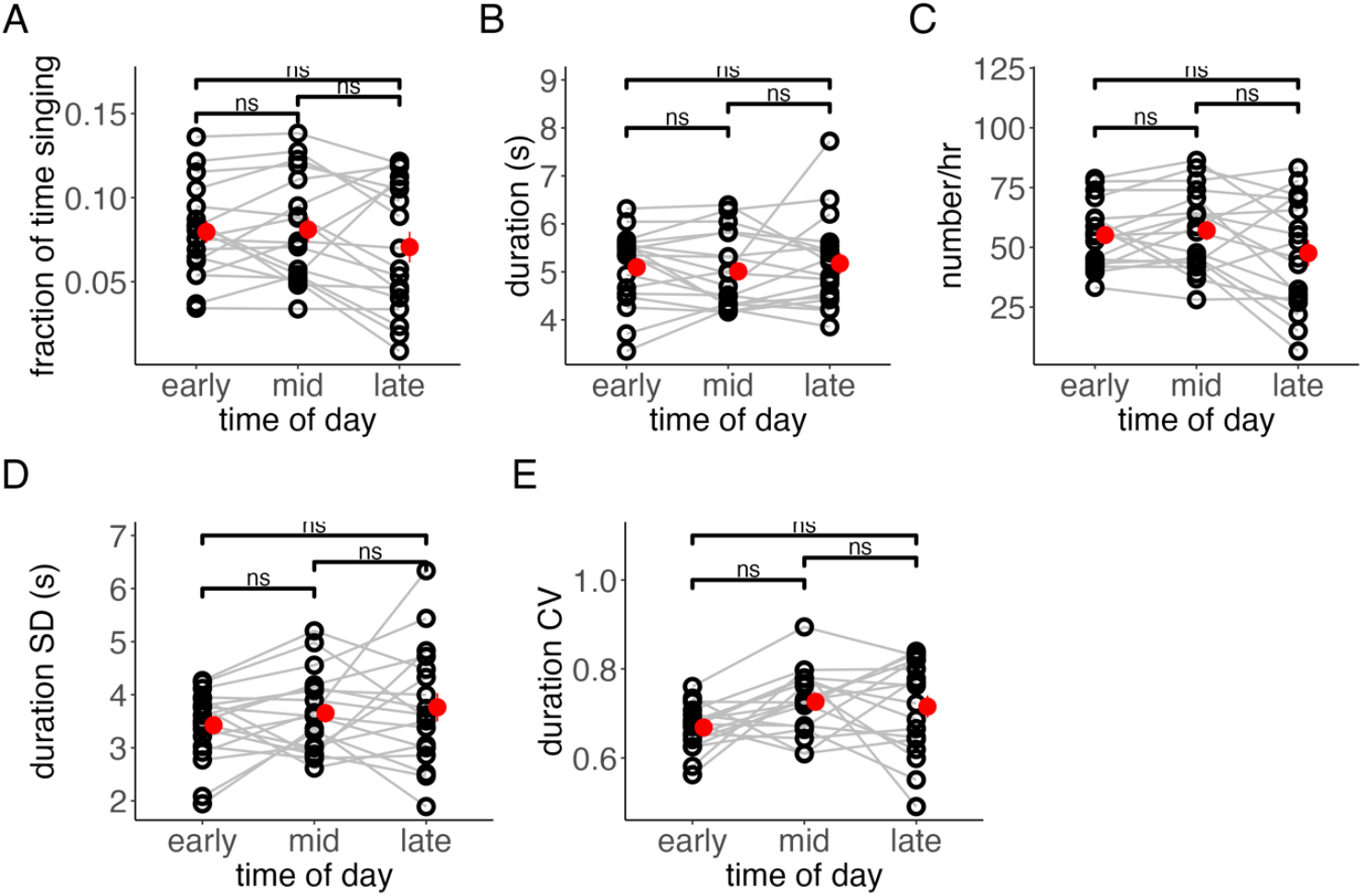
Daily variation of solo songs in a group of canaries. **A**. Fraction of time singing. **B**. Mean song duration. **C**. Number of songs per hour. **D**. Standard deviation (SD) of song duration. **E**. Coefficient of variation (CV) of song duration. Open black circles in each plots represent a day. The red filled circles indicate the mean and the red bar the standard error. Early, mid and late represent the interval times during the day, calculated by dividing each day into three equal parts. ns, p>0.05. Wilcoxon rank-sim test with Bonferroni-Holm correction.

**Supplementary figure S2.**
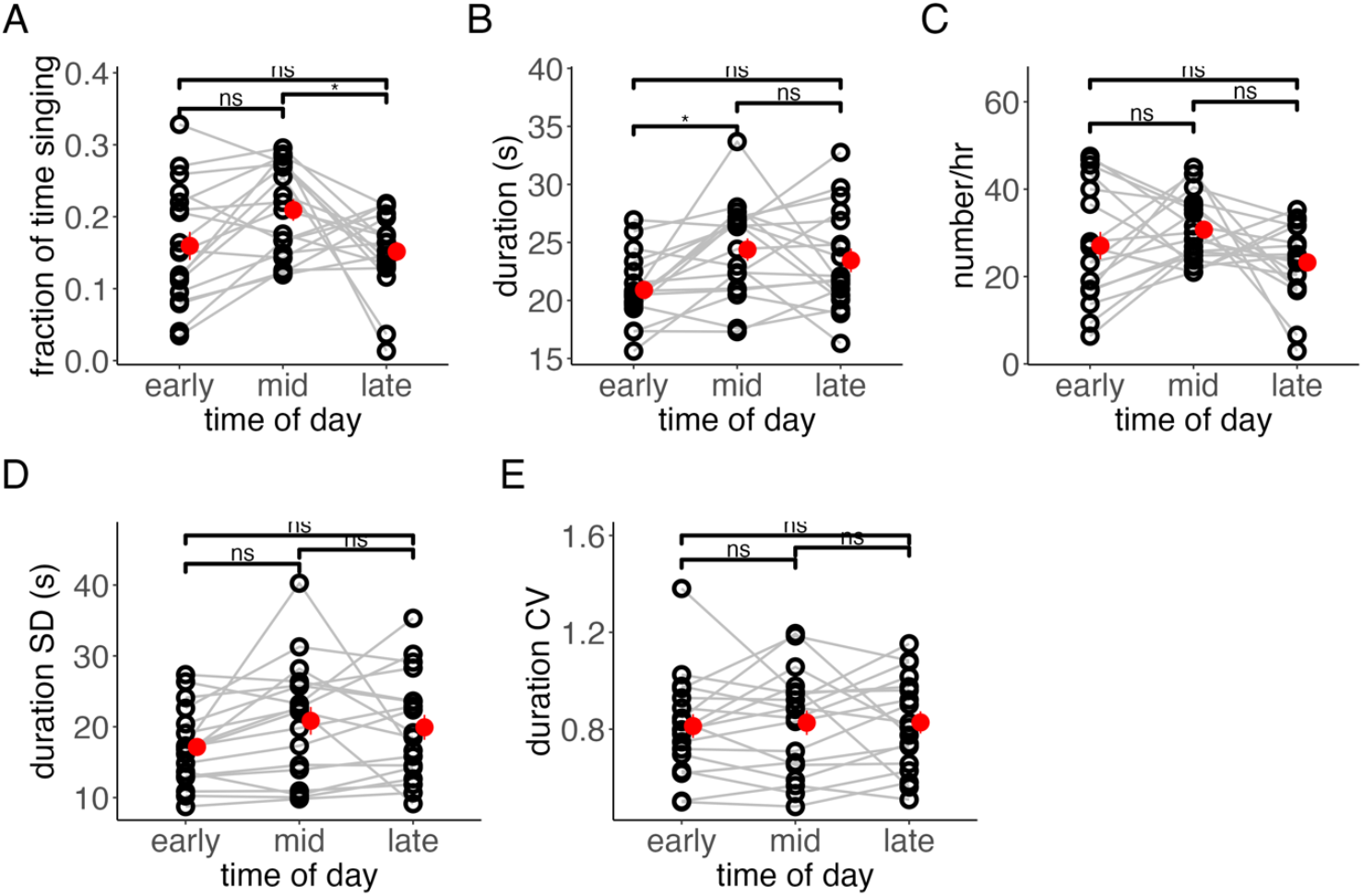
Daily variation of overlapping songs in a group of canaries. **A**. Fraction of time singing. **B**. Mean song duration. **C**. Number of songs per hour **D**. Standard deviation (SD) of song duration. **E**. Coefficient of variation (CV) of song duration. Open black circles in each plot represent a day. The red filled circles indicate the mean and the red bar the standard error. Early, mid and late represent the interval times during the day, calculated by dividing each day into three equal parts. *, p<0.05; ns, p>0.05; paired Wilcoxon rank-sim test with Bonferroni-Holm correction.

